# Poly (acetyl, arginyl) glucosamine disrupts *Pseudomonas aeruginosa* biofilms and enhances bacterial clearance in a rat lung infection model

**DOI:** 10.1101/2021.09.15.460521

**Authors:** Bryan A. Garcia, Melissa S. McDaniel, Allister J. Loughran, J. Dixon Johns, Vidya Narayanaswamy, Courtney Fernandez Petty, Susan E. Birket, Shenda M. Baker, Roxana Barnaby, Bruce A. Stanton, Jeremy B. Foote, Steven M. Rowe, W. Edward Swords

## Abstract

*Pseudomonas aeruginosa* is a common opportunistic pathogen that can cause chronic infections in multiple disease states, including respiratory infections in patients with cystic fibrosis (CF) and non-CF bronchiectasis. Like many opportunists, *P. aeruginosa* forms multicellular biofilm communities that are widely thought to be an important determinant of bacterial persistence and resistance to antimicrobials and host immune effectors during chronic/recurrent infections. Poly (acetyl, arginyl) glucosamine (PAAG) is a glycopolymer which has antimicrobial activity against a broad range of bacterial species, and also has mucolytic activity which can normalize rheologic properties of cystic fibrosis mucus. In this study, we sought to evaluate the effect of PAAG on *P. aeruginosa* bacteria within biofilms *in vitro*, and in the context of experimental pulmonary infection in a rodent infection model. PAAG treatment caused significant bactericidal activity against *P. aeruginosa* biofilms, and a reduction in the total biomass of preformed *P. aeruginosa* biofilms on abiotic surfaces, as well as on the surface of immortalized cystic fibrosis human bronchial epithelial cells. Studies of membrane integrity indicated that PAAG causes changes to *P. aeruginosa* cell morphology and dysregulates membrane polarity. PAAG treatment reduced infection and consequent tissue inflammation in experimental *P. aeruginosa* rat infections. Based on these findings we conclude that PAAG represents a novel means to combat *P. aeruginosa* infection, which may warrant further evaluation as a therapeutic.

## Introduction

*Pseudomonas aeruginosa* is a Gram-negative bacterium that is frequently implicated in respiratory infections, neutropenic sepsis, and hospital acquired infections (1-4*). P. aeruginosa* is commonly isolated from the airways of patients with cystic fibrosis (CF) and other airway diseases impacted by chronic infection, including non-CF bronchiectasis and COPD (5, 6). Given the growing global incidence of antibiotic resistant infections by *P. aeruginosa*, the World Health Organization recently classified this species as a critical priority for the research and development of new antimicrobials (7).

In the lung, *P. aeruginosa* is a particularly difficult pathogen to treat due to its propensity to form biofilms, an adaptation that allows *P. aeruginosa* persist in spite of antimicrobial therapy and immune effectors (1, 8-10). Formation of airway biofilms by *P. aeruginosa* is a critical step in the development of chronic colonization of the bronchiectatic airway and is associated with worsened prognosis and increased utilization of antibiotics (11, 12). Biofilms are highly complex 3-dimensional ecosystems and less than 10% of airway biofilm biomass is bacterium (10, 13). The primary components of *P. aeruginosa* biofilm biomass are extracellular polymeric substances including polysaccharides and alginate that are secreted by *P. aeruginosa* (14). In airway biofilms, additional components from the local airway environment are also involved, including gel forming mucins and cellular debris such as extracellular DNA (13, 15, 16). Together, these substances form a gel-like matrix scaffold which provides survival of *P. aeruginosa* and other airway pathogens within biofilm communities (14).

Recently, a polycationic glycopolymer, poly (acetyl, arginyl) glucosamine (PAAG, now being developed as SNSP113) has been shown to improve mucus viscoelasticity and mucociliary clearance in CF rats and ferrets (17). In patients with CF, the architecture of the gel-forming mucin MUC5B is distorted due to deficient bicarbonate and an acidic pH that prevent calcium chelation and removal from the anionic mucin (17-19). This excessive Ca^2+^ in the MUC5B limits normal mucus expansion and results in a more globular and viscous form of the protein that is intrinsically more adherent and less amenable for mucociliary transport (19, 20). PAAG interferes with Ca^2+^ binding to MUC5B, restoring normal MUC5B architecture and improving its viscoelastic properties which in turn, enhance the function of the mucociliary clearance apparatus *in vitro* and *in vivo* (17). There are known roles for calcium in maintenance of the integrity of the bacterial outer membrane (21, 22), as well as in bacterial attachment to surfaces (23) and formation of biofilms (24). Studies have shown that calcium chelation disperses biofilms *in vitro* (25). Supporting this notion, treatment with PAAG significantly diminished biofilm density for *Burkholderia cepacia* complex (Bcc (26, 27). Given these previous findings, in this study we sought to identify anti-microbial and anti-biofilm efficacy of PAAG against *P. aeruginosa* using laboratory and clinical isolates *in vitro* biofilm models, and respiratory infections *in vivo*.

## Methods

### Bacterial Strains and Growth Conditions

Bacterial strains of *P. aeruginosa* utilized included a well characterized non-mucoid *P. aeruginosa* strain (PAO1; ATCC#BAA-47) and *P. aeruginosa* PA529,a mucoid clinical isolate obtained from an adult CF patient at the UAB Cystic Fibrosis clinic. Bacterial strains were stored at −80°C in tryptic soy broth (TSB) or Mueller-Hinton broth (MHB) containing 15% (vol/vol) glycerol. All strains of *P. aeruginosa* were routinely cultured on tryptic soy agar (TSA) or in tryptic soy broth (TSB) unless otherwise stated. *P. aeruginosa* strains were streaked for colony isolation before inoculating into LB broth and shaking overnight at 37°C, 200 rpm.

### Minimum biofilm eradication concentration (MBEC)

To test the minimum biofilm eradication concentration (MBEC) of PAAG *against P. aeruginosa* biofilms, overnight bacterial cultures were cultured to logarithmic phase of growth, and diluted to an approximate OD_600_ of 0.25 (∼10^8^ cells/mL) and added to wells of a 96-well microtiter dish. An MBEC™ High-throughput (HTP) Assay (Innovotech) lid with sterile peg inserts was added and the inoculated plates were incubated for 48 hours at 37°C on a shaker (3-5 rpm/min). Pegs were then rinsed twice with PBS and placed in a 96-well flat bottom microtiter plate containing 50-200 µg/ml of PAAG formulated in sterile water and incubated at room temperature for 1 hour. The plate lid was then transferred to a fresh plate containing PBS and sonicated to remove adherent bacteria. To calculate the MBEC, biofilms were serially diluted, plated onto MHB agar plates for viable colony counting.

### CSLM imaging and biofilm quantification

Confocal laser scanning microscopy (CLSM) was performed using a Nikon-A1R HD25 Confocal Laser Microscope (Nikon, Tokyo, Japan). Images were acquired and processed using the NIS-elements 5.0 software. For the purpose of confocal imaging, *P. aeruginosa* PAO1 expressing the GFP-containing plasmid pSMC21 (28) was grown overnight in TSB and diluted to an OD_600_ of 0.075 (∼10^7^ cells/mL). *P. aeruginosa* were then cultured in TSB (0.3 mL) in 35-mm glass bottom well confocal petri dishes (MatTek corp. Ashland, MA, USA) and biofilms were allowed to grow for 24 hours at 37°C in a humidified 5% CO_2_ incubator. Biofilms were treated with 0.1 mL vehicle (PBS) or PAAG for 3 hours before imaging. At least five biological replicates were imaged for each condition, with the mean maximal biofilm thickness of three technical replicates used for statistical analysis. All representative images were projected using alpha-blending. Quantitative measures of biofilm structure were obtained from confocal Z-stack images using COMSTAT image analysis software.

### CSLM imaging of LIVE/DEAD stained biofilms

To assess the impact of PAAG on the structure of mature *P. aeruginosa* biofilms, *P. aeruginosa* PAO1 was prepared as a broth culture in TSB for 24 hours and then diluted to an OD_600_ of 0.075 (∼10^7^ cells/mL). Confocal dishes were inoculated with 0.3 mL of diluted culture, and biofilms were incubated at 37°C in a humidified 5% CO_2_ incubator for 24 hours. Biofilms were then treated for 3 hours with vehicle or PAAG, essentially as described above. After the treatment period, LIVE/DEAD staining was performed using LIVE/DEAD™ *Bac*Light™ Bacterial Viability Kit. Representative images were displayed using maximum intensity projection.

*P. aeruginosa* immortalized bronchial epithelial cells for 6 hours. PAAG was prepared at a concentration of 250 ug/ml and added at concentration as indicated.

### Influence of calcium on PAAG anti-biofilm activity

The experiment evaluated PAAG’s antibiofilm activity on *P. aeruginosa* biofilms in a calcium rich environment. Biofilms were grown from an overnight culture of *P. aeruginosa* PAO1 or *P. aeruginosa* PA529 in TSB. Overnight cultures were adjusted to an approximate OD_600_ of 0.25 (∼10^8^ cells/mL), and then diluted 1:30 in TSB. The diluted culture (0.1 mL) was then added to each well of a 96-well microtiter dish in triplicate. Bacteria was grown in a stationary incubator for 48 hours, before washing with 1X PBS to remove planktonic bacteria. Then the biofilms were treated with PAAG 2-16 (100-500 ug/ml) alone or in combination with CaCl_2_ (200 nM) and incubated at room temperature for 24 h. Biofilms treated with media was used as an untreated control. After 24 h, the wells were rinsed twice with PBS and dried for 2 h at 37°C. The biofilms were stained with 1% crystal violet for 15 min. Stained biofilms were washed twice with PBS and crystal violet was eluted in absolute methanol. After incubation for 5 min, the solubilized crystal violet was transferred into a fresh microtiter plate and the optical density (OD) was read at 590 nm. The average absorbance of biofilm-forming isolates was greater than the average absorbance of the negative control wells +/-3 standard deviations, confirmed biofilm formation.

### Cell adherence after pretreatment with PAAG

Planktonic cultures *of P. aeruginosa* were grown in TSB for 24 h at 37°C in a shaking incubator. Cultures were then reseeded and grown to mid-logarithmic phase (OD_600_ ∼= 0.6), washed in PBS and diluted to ∼10^6^ CFU/ml in F12K media. A549 immortalized human type II pneumocyte (ATCC#CCL-185) cells were grown as monolayers to 85-95% confluence in 24-well tissue culture plates. The number of eukaryotic cells per well was determined by quantitation via hemocytometer; the average number of cells per well was 50,000 cells/well. PAAG (50-500 ug/mL) diluted in F12K medium or a medium-only control was added to each well and plates were incubated for 1 hour at 37°C with 5% CO_2_. Wells were then washed, and each well was inoculated with diluted bacterial culture. The infected monolayers were centrifuged (165 x g for 5 min) and incubated at 37°C with 5% CO_2_ for 1 hour, then washed to remove any nonadherent bacteria. The wells were trypsinized (5 min) and lysed with 0.25% Triton X-100 (5 min). Cell layers were removed from the surface by scraping before being serially diluted and spot plated to quantify viable bacteria. All assays were performed in triplicate.

### Cytoplasmic Membrane Depolarization Assay

Membrane depolarization of *P. aeruginosa* was measured using the membrane potential-sensitive fluorescent dye 3,3′-Dipropylthiadicarbocyanine iodide (DiSC3-5) (29). Overnight cultures of *P. aeruginosa* strains PAO1 and PA529 were pelleted by centrifugation (13,000 rpm for 1 min) and washed three times with 5 mM HEPES (pH 7.8) buffer. Cells were then resuspended in 5 mM HEPES (pH 7.8) buffer with 0.2 mM EDTA to an OD_600_ of 0.05 (45). DiSC3(5) and KCl were added to a final concentration of 0.4 µM and 0.1 M, respectively, and cultures were shaken (37°C, 150 rpm) for 20 min until fluorescence quenching was achieved. PAAG was added at a variety of concentrations (50–500 μg/mL) and fluorescence was measured at an excitation wavelength of 622 nm and an emission wavelength of 670 nm after 15 min of incubation. Treatment with triton X-100 (0.1% v/v) was used as the positive control for membrane depolarization. The experiment was performed in triplicate with three independent cultures.

### Propidium Iodide (PI) Uptake Assay

Overnight cultures of *P. aeruginosa* were pelleted by centrifugation (13,000 rpm for 1 min) and supernatants were discarded before resuspending in water to an OD_600_ of 0.25 (∼10^8^ cells/mL). PI was used at a concentration of 17 μg/mL as described in previous studies (26). All assays were performed at room temperature. PI was added to the wells of a 96-well plate containing ∼10^8^ CFU/mL of bacterial suspension. Fluorescence measured via SpectraMax Gemini XPS (Molecular Devices). Increasing concentrations of PAAG (50, 100, 250 and 500 μg/mL) were added to the wells. Cells treated with 0.1% Triton X-100 were used as a positive control for membrane permeability. PI alone and PI on untreated cells were included as negative controls. Fluorescence was measured with an excitation wavelength of 535 nm and emission wavelength of 625nm every 10 minutes for 4 h. The experiment was performed in triplicate with three independent cultures.

### Scanning electron microscopy

For scanning electron microscopy (SEM) imaging, planktonic cells of *P. aeruginosa* PAO1 or *P. aeruginosa*PA529 were grown for 24 hours on silicon wafers placed in sterile 12-well plates. PAAG (250 µg/mL) was added and the plate was incubated for 1 hour. The planktonic cells were fixed overnight with 2.5 % glutaraldehyde and 0.5% paraformaldehyde in 0.1 M phosphate buffer and then rinsed with 0.1 M phosphate buffer (3 × 10 min/wash). The planktonic cells were then dehydrated gradually via sequential washes with 10, 25, 50, 75, and 95 % alcohol (5 min/wash) and 100 % alcohol (3 × 5 min/wash). Hexamethyldisilane (HMDS) was used for overnight drying. Samples were then sputter-coated with gold prior to SEM imaging.

*Prevention of in-vivo P. aeruginosa infection via pre-treatment with nebulized PAAG* Sprague Dawley rats (*Rattus norvegicus*, age 6 months) were administered nebulized PAAG (250 μg/ml x 20 ml over 45 min) in isotonic glycerol or glycerol vehicle control on days 1, 3, 5, 7. On day 7 post treatment, rats were infected intratracheally with *P. aeruginosa* (∼10^5^ CFU/animal). Rats continued to receive nebulized PAAG or control for two doses at day 10 and 12 post-infection at which time they were euthanized. The left lung of each mouse was harvested and homogenized in DMEM (Gibco) for viable plate counting. Homogenate was plated on PIA and grown 24 hours at 37°C to obtain viable CFU counts. The right lung of each animal was inflated with 10% buffered formalin and stored at 4°C for histological analysis. All animal procedures were performed using standard procedures according to AVMA guidelines and were fully reviewed and approved by the UAB Institutional Animal Care and Use Committee (IACUC).

### Histological analysis

Inflated right lungs from infected rats were stored in 10% neutral buffered formalin (Fisher Scientific, Waltham, MA) at 4°C until processing. Sections from each lobe of the right lung were trimmed and sent to the UAB Comparative Pathology Laboratory to be paraffin embedded and hematoxylin and eosin (H&E) or periodic acid-Schiff (AB-PAS) stained. Images of stained lung sections were taken on a Leica LMD6 scope (Wetzlar, Germany) at 10 x magnification. Semi-quantitative grading of all lung sections was performed by a board-certified veterinary pathologist in a blinded fashion (J.F.). For the H&E-stained sections, semi-quantitative histopathological scores were assigned using a scoring matrix based primarily on neutrophilic influx. Severity was rated on a scale of 0-4 where 0 represents no observable neutrophils, 1 for ≤25%, 2 for 25-50%, 3 for 50-75%, or 4 for 75-100% of the section affected of independently viewed fields of view. Density of PMNs ranged in affected fields was scored from 1-3, to represent mild, moderate, and severe respectively. The final histopathological score was calculated by multiplication of extent and severity scores.

### Statistical analysis

Unless otherwise noted, graphs represent sample means ± SEM. Differences between group were analyzed via one-way ANOVA assuming a Gaussian distribution with Dunnett’s multiple comparison tests unless otherwise stated. *P*-values less than 0.05 were considered statistically significant. Statistical analysis was performed using Graphpad Prism Version 6.0 (San Diego, CA).

## Results

### PAAG treatment disrupts biofilm structure

PAAG has previously been demonstrated to have bactericidal effects on planktonic *Pseudomonas aeruginosa, Staphylococcus aureus* and *Burkholderia cepacia* (26, 27). However, we wanted to address the effect of PAAG on pre-formed, mature *P. aeruginosa* biofilms. To determine the anti-biofilm activity of PAAG, biofilms of *P. aeruginosa* PAO1, and a mucoid clinical isolate, *P. aeruginosa* PA529, were allowed to grow and mature for 48 hours and were subsequently exposed to varying concentrations of PAAG (0-500 μg/ml) for one hour using a standard MBEC assay. Exposure of mature biofilms to PAAG at 50 µg/ml or above resulted in a statistically significant reduction in viable bacterial counts for both *P. aeruginosa* PAO1 and *P. aeruginosa* PA529 (*P*<0.0001) (Figure 1A,B). The effect of PAAG on preformed *P. aeruginosa* PAO1 (GFP+) biofilms was visualized using CSLM imaging. Exposure of the *P. aeruginosa* PAO1 biofilm to PAAG treatment resulted in biofilm reduction, as seen in representative confocal images (Figure 1 C,D). When quantified, there was a significant reduction in mean maximal biofilm thickness from (25.3 μm ± 4.9 μm for control versus 8.5 μm ± 3.5 μm for PAAG, *P*<0.01), and in total biomass (0.005 μm^3^/μm^2^ ± 0.003 μm^3^/μm^2^ for control versus 0.002 μm^3^/μm^2^ ± 0.0009 μm^3^/μm^2^ for PAAG, *P*<0.05). The surface area of PAAG treated biofilms was also decreased, albeit below levels of statistical significance (3381 μm^2^ ± 1595 μm^2^ for control versus 1494 μm^2^ ± 1.003 μm^2^ for PAAG, *P*=0.0556) (Figure 1E). These findings demonstrate that PAAG antibacterial efficacy against *P. aeruginosa* is preserved even in the setting of a well matured biofilm, and that PAAG induces destabilization of the usual biofilm architecture.

**Figure 1.**
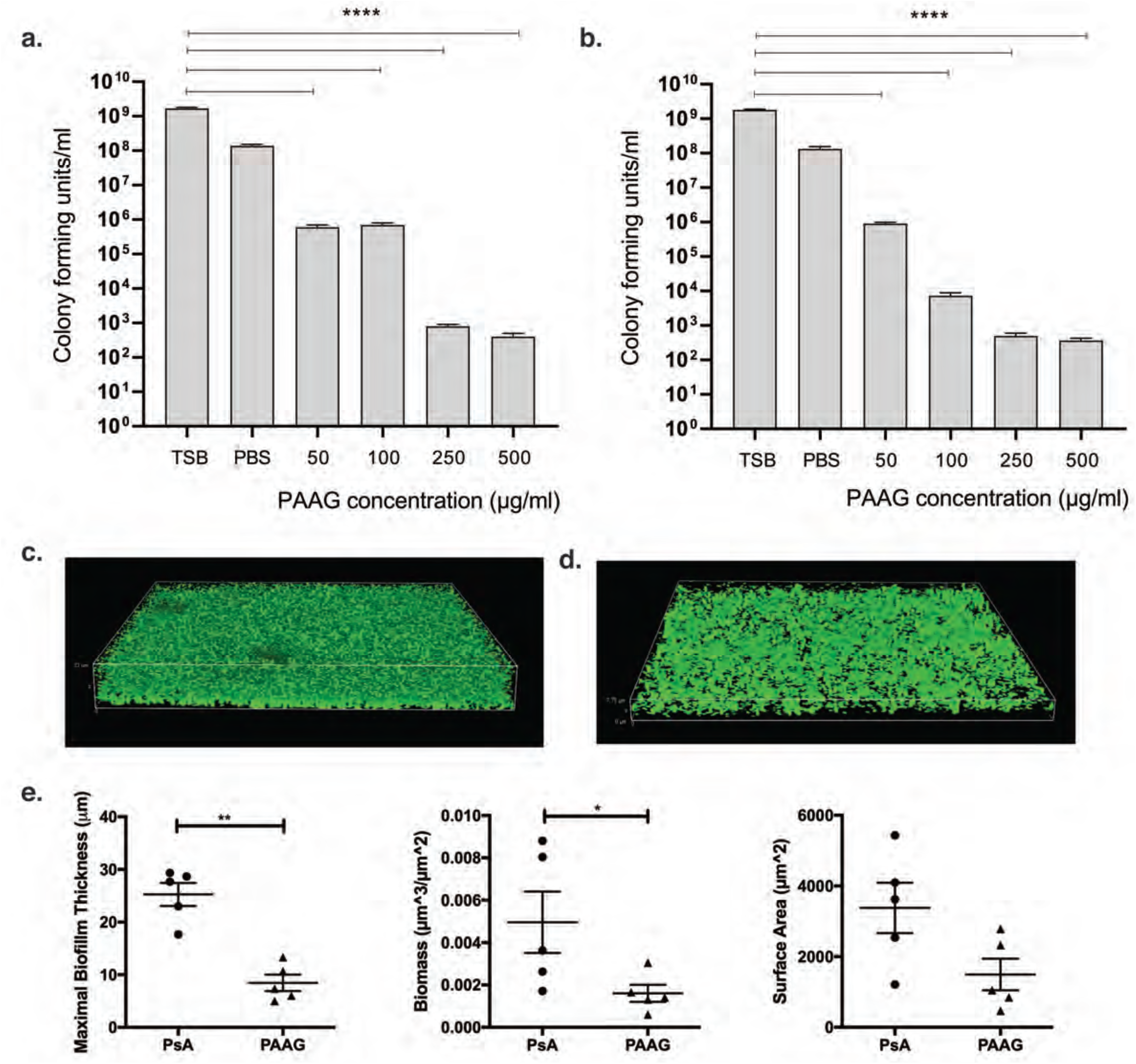
PAAG shows antimicrobial and antibiofilm effects against *P. aeruginosa* biofilms. Biofilms of A) *P. aeruginosa* PAO1 and B) *P. aeruginosa* PA529 were grown for 48 h on minimum biofilm eradication concentration (MBEC) plates before exposure to various concentrations of PAAG for 1 hour. Viable cells were quantified as colony forming units/mL. **** P<0.0001. C,D) Effect of PAAG (200 μg/ml) as compared to untreated controls evaluated by confocal microscopy. E) Maximal biofilm thickness, biomass, and surface area of biofilms were measured via COMSTAT software. *P<0.05, **P<0.01.

### PAAG shows antimicrobial effects of on intact *P. aeruginosa* biofilms

Given our findings of antimicrobial activity of PAAG on planktonic *P. aeruginosa* and the ability of PAAG to disrupt the structural properties of *P. aeruginosa* biofilms, we next assessed were visualized antimicrobial effect of PAAG on a preformed biofilm of *P. aeruginosa* PAO1 using BacLight LIVE/DEAD (Molecular Probes) staining and CSLM imaging. Syto9 stained the viable bacteria (green) throughout the biofilm treated with PBS control (Figure 2A). Treatment with PAAG resulted in significant bacterial killing throughout all levels of the biofilm including at the deepest layers where the biofilm and glass-substratum interfaced (Figure 2B).

**Figure 2.**
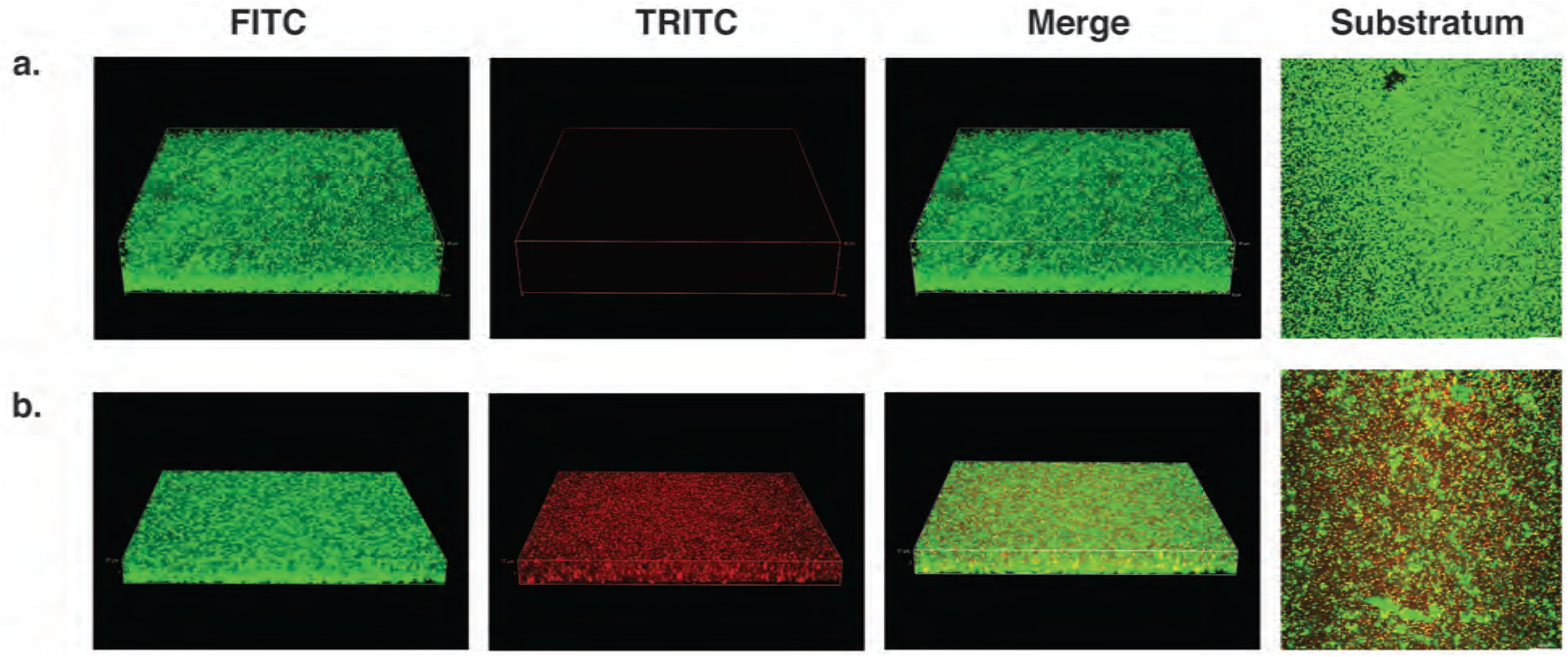
Antimicrobial effect of PAAG on *P. aeruginosa* PAO1 biofilms by live/dead imaging. *P. aeruginosa* PAO1 biofilms treated with A) vehicle control or B) PAAG (250 μg/ml) were visualized by confocal imaging after live/dead staining. Single channels of live bacteria (green) and dead bacteria (red) are represented, in addition to a merged image from a single z-series acquisition along with a representative image of the substratum layer.

### PAAG treatment decreases *P. aeruginosa* biofilm formation on bronchoepithelial cells

PAAG significantly decreased *P. aeruginosa* biofilm and showed antimicrobial effects on bacteria grown on abiotic surfaces. However, the efficacy of PAAG in the disease-relevant context of bacterial biofilms grown on a respiratory epithelium has not yet been evaluated. Biofilms of *P. aeruginosa* PAO1 (GFP+) were cultured on confluent monolayers of CFTR -/- cystic fibrosis bronchoepithelial cells (CFBEs) for 6 hours before treating with PAAG (250 µg/mL). Biofilm formation was visualized via CLSM imaging and quantified using COMSTAT. Exposure of *P. aeruginosa* PAO1 biofilms grown on CFBEs to PAAG treatment again resulted in biofilm reduction, as seen in representative confocal images (Figure 3 A,B). Quantification of biofilm indicated a significant reduction in mean maximal biofilm thickness (32.7μm ± 4.7 for control vs 12.2μm ± 4.1 for PAAG, *P*<0.01), in total biomass (0.008 μm^3^/μm^2^ ± 0.002 μm^3^/μm^2^ for control versus 0.003 μm^3^/μm^2^ ± 0.001μm^3^/μm^2^ for PAAG, *P*<0.01) and in surface area of the biofilm (3630 μm^2^ ± 995 μm^2^ for control versus 1694 μm^2^ ± 700.5 μm^2^ for PAAG, *P*<0.05) (Figure 3C). These findings demonstrate that the ability of PAAG to disrupt and permeabilize *Pseudomonas* biofilms is retained in the clinically relevant context of coculture with an epithelial cell layer.

**Figure 3.**
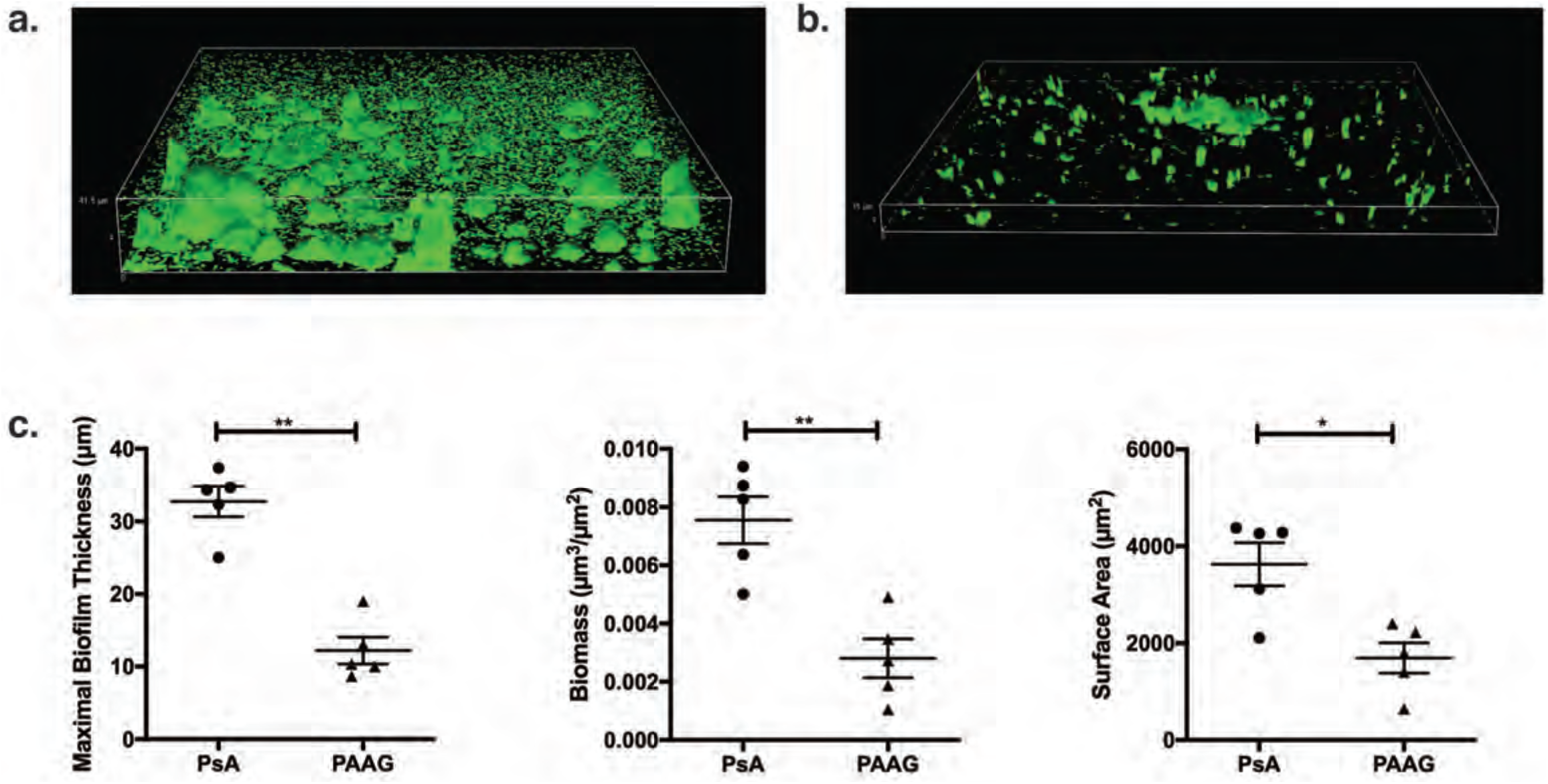
PAAG maintains antibiofilm effects against biofilms on epithelia. Biofilms of *P. aeruginosa* PAO1 were grown on confluent monolayers of CFTR -/- bronchoepithelial cells (CFBE) for 6 hours before a 1 hour treatment with PAAG (250 μg/ml). **** P<0.0001. The effect of PAAG was evaluated via confocal imaging of A) untreated and B) PAAG treated biofilms. C) Changes in biofilm structure including maximal biofilm thickness,, biomass, and surface area of biofilms were measured via COMSTAT software. *P<0.05, **P<0.01.

### Calcium chloride abrogates the effect of PAAG *on P. aeruginosa* biofilm dissolution

*P. aeruginosa* and other gram-negative bacteria have their outer membrane coated with polyanionic lipopolysaccharide molecules (LPS) that provide structural integrity to the membrane of the bacteria by cross linking with divalent cations including calcium (30). There is also precedent for calcium playing similar crosslinking roles for extracellular DNA or polysaccharide in biofilm matrix (23, 24). Since our previous experiments demonstrated a potent biofilm removal effect by PAAG and because PAAG has previously been shown to displace calcium from mucins (17), we next sought to assess if PAAG similarly causes calcium displacement to achieve biofilm dissolution. *P. aeruginosa* PAO1 or *P. aeruginosa* PA529 biofilms were treated with PAAG (100-500 µg/mL) with and without added CaCl_2_ (200nM). Remaining biofilm was quantified by crystal violet staining (absorbance at OD_590_). PAO1 biofilms exposed to PAAG in a calcium rich environment required a PAAG concentration of 500µg/mL to decrease biomass (OD_590_=0.89 ± 0.07 vs 0.38 ± 0.07, *P<*0.01) whereas a PAAG concentration of 100 µg/mL is sufficient without excess calcium (OD_590_=0.89 ± 0.07 vs 0.33 ± 0.02, *P<*0.0001) (Figure 4A). For biofilms of *P. aeruginosa* PA529, the addition of calcium abrogated the effects of PAAG at all concentrations tested (Figure 4B). These findings suggest that dissolution of intact biofilms by PAAG may occur as a result of calcium displacement, which could either impact the stability of the biofilm matrix or the viability of bacteria present in the biofilm.

**Figure 4.**
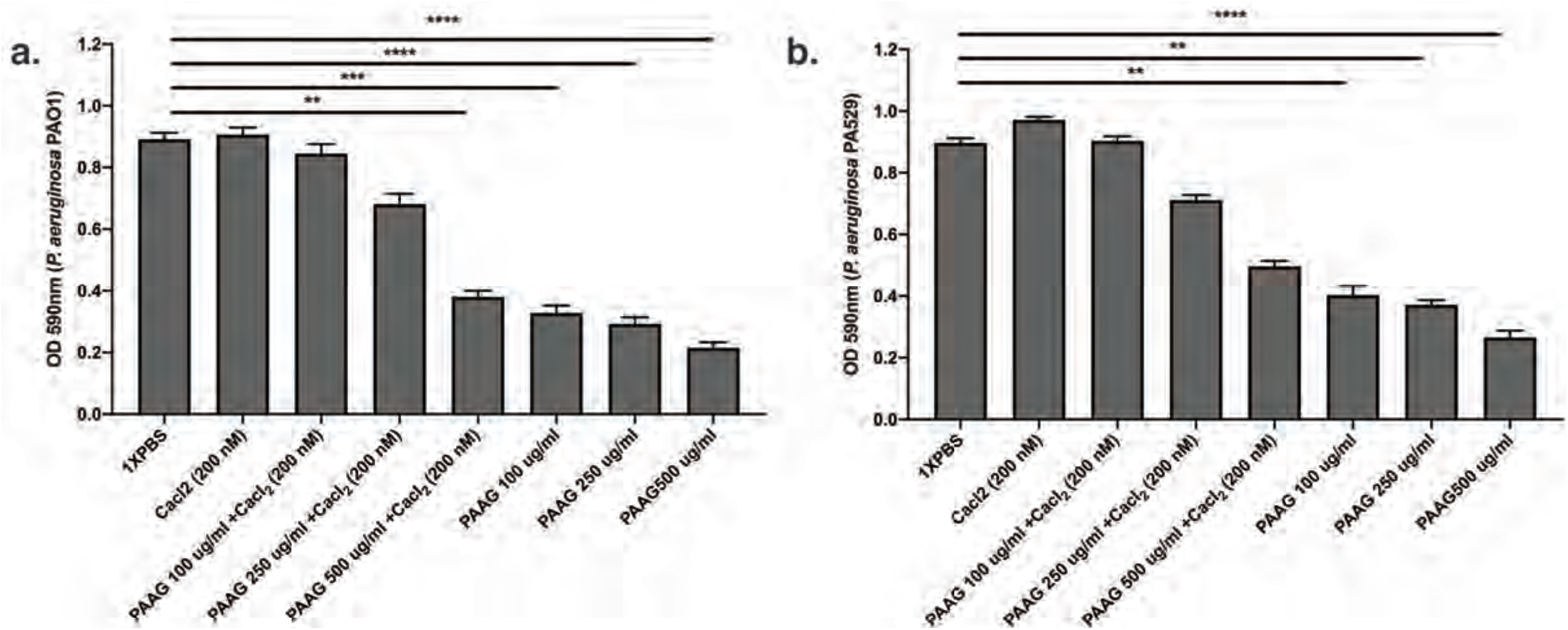
Biofilm dissolution by PAAG is abrogated by Ca^2+^. *P. aeruginosa* SUS116 biofilms were exposed to 200 μg/ml of PAAG and either sterile water or calcium chloride. Co-exposure of the biofilms to PAAG and calcium chloride resulted in reduced biofilm dissolution compared to co-exposure with sterile water (87.2% reduction in sterile water vs. 58.9% reduction in 150 mM calcium chloride, P<0.0001).

**Figure 5.**
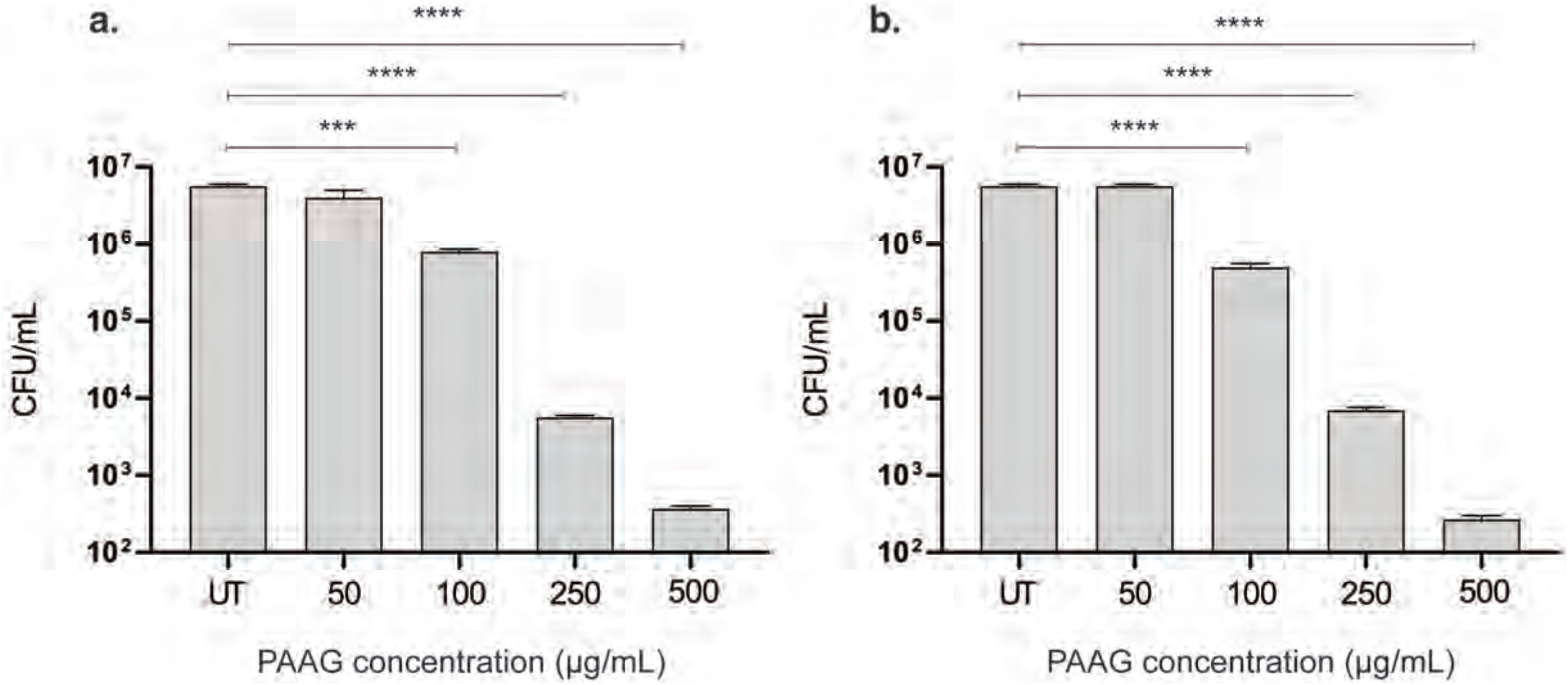
Pretreatment with PAAG reduced adherence of P. aeruginosa to A549 cells. A549 immortalized lung carcinoma cells were pretreated for 1 h with PAAG (0-500 μg/ml), washed, and then incubated with ∼10^6^ cells *P. aeruginosa* PAO1 **(A)** or *P. aeruginosa* Pa529 **(B)**. Subsequently, to estimate the number of bacteria attached, A549 cells were exposed to detergent Triton X-100, washed, and the lysate serially diluted in sterile PBS and plated onto tryptic soy agar (TSA) plates to determine the number of CFU per well. Untreated A549 cells incubated with ∼10^6^ *P. aeruginosa* PAO1 or *P. aeruginosa* Pa529. **** P<0.0001, *** P=0.0001, compared to the untreated control of the respective strains.

### PAAG prevents *P. aeruginosa* bacterial adhesion on A549 cell surface

Calcium displacement is known to interfere with adherence of *P. aeruginosa* to respiratory epithelia (31). Given the findings that PAAG treatment resulted in bacterial killing at the substratum, we next sought to determine if PAAG altered *P. aeruginosa* attachment to immortalized lung pneumocytes (A549) using a cell adherence assay. Both *P. aeruginosa* PAO1 and *P. aeruginosa* PA529 adhered to A549 cells treated with vehicle/ untreated control (UT). However, pretreatment with PAAG resulted in a dose-dependent reduction decrease in the adherence of *P. aeruginosa* (Figure 7A,B). PAAG at a concentration of 250 ug/ml resulted in 3 -log reduction in the number of bacteria on the surface of A549 cells for both strains of *P. aeruginosa* tested (P<0.0001).

### PAAG exposure results in bacterial membrane depolarization and permeability

Calcium displacement has been shown to compromise integrity of the bacterial membrane. We hypothesized that this might be mechanism by which PAAG produces antimicrobial effects on *P. aeruginosa* (21). DiSC3, a membrane potential-sensitive dye was used to evaluate the depolarization potential of PAAG on *P. aeruginosa* membranes. PAAG treatment resulted a rapid increase in DiSC3 fluorescence, reflecting depolarization of the electrical potential gradient (Figure 6 A,B). Exposure to PAAG at concentrations equal to or greater than their MIC, lead to rapid permeabilization of the cytoplasmic membranes in both strains of *P. aeruginosa*, thus depolarizing the electric potential gradient resulting in release of diSC3 and a consequent increase in fluorescence. Outer membrane permeabilization of the *P. aeruginosa* isolates was determined via propidium iodide (PI) uptake assay. PAAG also rapidly permeabilized the outer membrane of both *Pseudomonas* isolates tested in a concentration-dependent manner, as indicated by an increase in PI fluorescence. Untreated cells demonstrated no change in PI fluorescence intensity (Figure 6 C,D).

**Figure 6.**
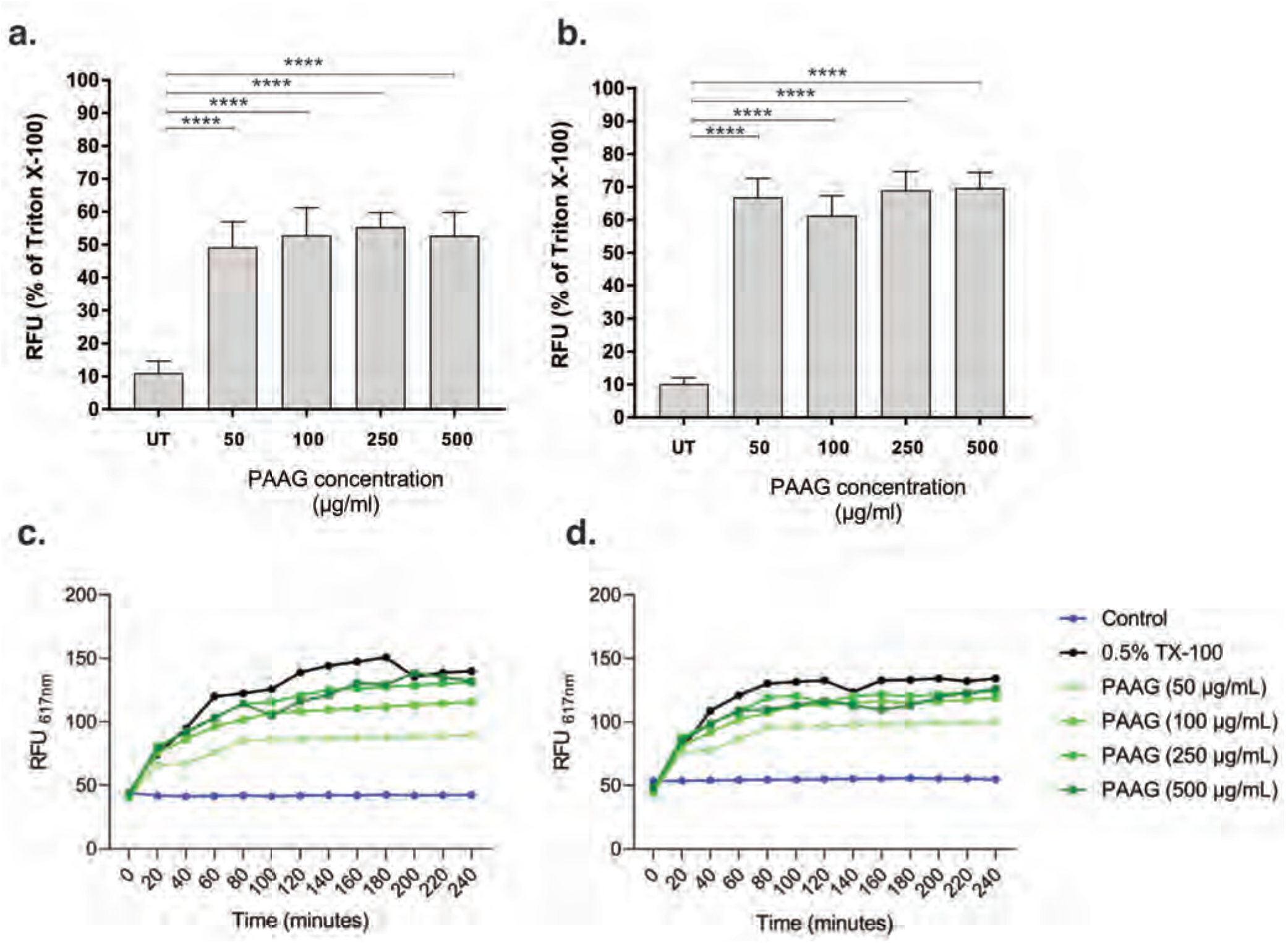
PAAG disrupts *P. aeruginosa* membrane integrity. Depolarization of the cytoplasmic membrane of A) PAO1 and B) PA529 by PAAG was measured using the membrane potential-sensitive dye, DiSC3. Results are represented as a percentage of the detergent Triton X-100 (TX-100) compared to the untreated control (UT) of the respective strains. **** P<0.0001. The ability of PAAG to disrupt the integrity of the outer membrane of C) *P. aeruginosa* PAO1 and D) *P. aeruginosa* PA529 was evaluated with the fluorescent dye propidium iodide (PI) and measured as relative fluorescent units (RFU). Changes in fluorescence were measured via spectrophotometer at 617 nm.

### PAAG exposure causes morphological changes in *P. aeruginosa*

. Following exposure to PAAG (250 μg/ml), morphologic changes were observed on the otherwise smooth outer bacterial surfaces of both PAO1 and PA529. The morphological changes are indicative of membrane disruption, leaving a jagged appearance on the surface, resulting in the collapse of the bacterial structure (Figure 7). This is consistent with rapid uptake of cell impermeable dye, propidium iodide observed in Figure 6.

**Figure 7.**
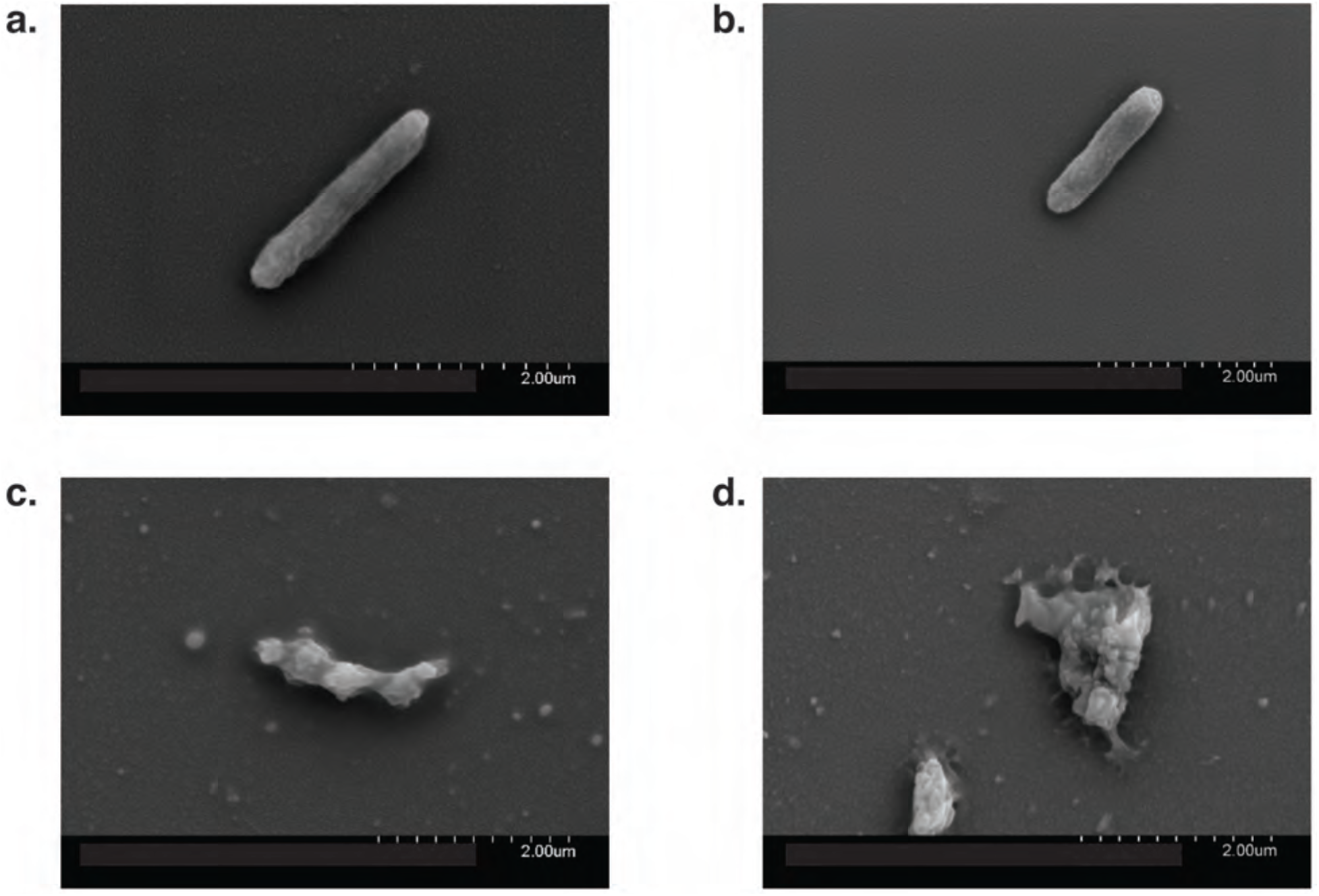
PAAG causes phenotypic changes to bacterial cell morphology. SEM images were taken of planktonic cultures of *P. aeruginosa* PAO1 (A, C) and *P. aeruginosa* PA529 (B, D); treated with vehicle control (A,B) or PAAG (C, D) for 60 minutes. Images representative of that observed in multiple independent samples.

### *In vivo* pre-treatment with PAAG decreases *P. aeruginosa* infection

Having demonstrated that PAAG has a bactericidal effect on biofilms of *P. aeruginosa* and reduces bacterial adhesion to an epithelial monolayer *in vitro*, we next sought *in vivo* confirmation of PAAG efficacy. We used a model of acute pulmonary infection by *P. aeruginosa* PA529 in Sprague-Dawley rats following pre-exposure to PAAG or vehicle control and concurrent treatment until euthanasia at 48 hours post infection (Figure 8A). This model sought to identify if PAAG can impair *P. aeruginosa* adhesion to an intact airway epithelium and reduce bacterial burden. Using this model, rats treated with PAAG demonstrated a significantly lower bacterial burden of *P. aeruginosa* in lung homogenate (2.38 log CFUs/mL in treated rats vs 4.86 log CFUs/mL in untreated controls, *P*<0.05) (Figure 8B). Lung histology revealed rats treated with control exhibited significant peri-bronchiolar inflammation and neutrophilic infiltrate following *P. aeruginosa* infection, whereas this finding was abrogated in the PAAG treatment group. Histopathologic analysis indicated significant inflammation in all the infected rats, with slightly diminished neutrophilic counts and peri-bronchial infiltration in rats treated with PAAG (Figure 8C).

**Figure 8.**
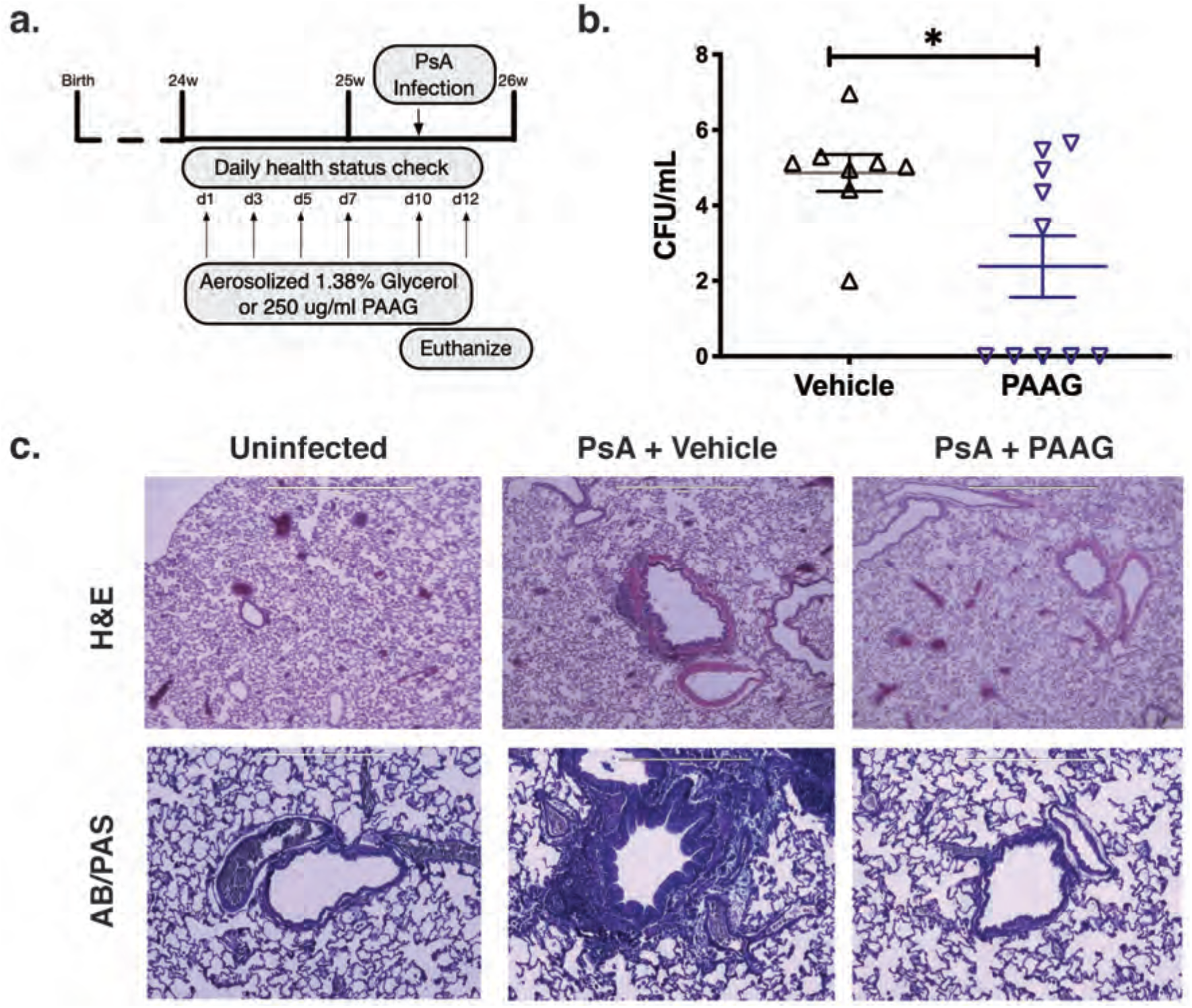
PAAG pre-treatment inhibits *Pseudomonas aeruginosa* infection *in vivo*. Sprague-Dawley rats were inoculated intratracheally with *P. aeruginosa* PA529 (∼10^5^ CFU), treated daily with nebulized PAAG (250 μg/ml x 20 ml over 45 min) or vehicle control (1% glycerol x 20 ml over 45 min) then euthanized 48 h later for analysis. **(A)**. Number of animals which grew *P. aeruginosa* (grey bars) or were sterile (black bars) *P < 0.05 by Fisher’s exact test. **(B)**. Bacterial load as measured by CFUs isolated post-treatment from vehicle control vs. PAAG-treated rats (∼ 10^6^ vs. ∼ 10^5^, *P < 0.05 by unpaired t-test). Dotted line represents initial inoculation level of 10^5^ CFUs. (**C)**. H&E (top) or AB-PAS staining of representative lung histology sections from animals without *P. aeruginosa* inoculation as compared to those with *P. aeruginosa* inoculation and either vehicle or PAAG treatment. Red arrow designates peribronchiolar neutrophilic inflammation and yellow arrow designates neutrophilic inflammation in the parenchyma. Less severe peribronchiolar and parenchymal neutrophilic inflammation was evident in PAAG-treated tissue. AB/PAS stained sections demonstrate reduced mucus expression in PAAG treated rats in the peribronchial region.

## Discussion

The global threat of rising antibiotic resistant infections has led to an emergent need to identify and develop new antimicrobials with novel and multimodal mechanisms of action (7). *P. aeruginosa* causes a wide variety of opportunistic infections that include respiratory infections in patients with CF. *P. aeruginosa* has a large and complex genome and rapidly develops antibiotic resistance against multiple classes of antibiotics making treatment difficult (3, 8, 11, 12, 32). *P. aeruginosa* persists within multicellular biofilm communities which are inherently resistant to antibiotics via multiple mechanisms including impaired antibiotic penetration, expression of antibiotic inactivating enzymes, reduced metabolic activity of biofilm imbedded bacteria, and augmented inter-bacterial genetic transfer (17, 18, 8, 12). Given the implications of biofilms impairing antibiotic efficacy, taking advantage of mechanisms that combat biofilm structural integrity and prevent bacterial biofilm formation is of substantial scientific and clinical importance.

The study highlights the bactericidal potential of PAAG against laboratory isolate *P. aeruginosa* within planktonic cultures or as a biofilm. The study also exemplifies its potential to disrupt and permeabilize the structure of mature biofilms formed by these isolates. PAAG has been studied and proven to be an effective bactericidal agent against planktonic cultures of *Pseudomonas aeruginosa, Staphylococcus aureus* and *Burkholderia cepacia* (26, 27). PAAG was also found to effectively remove pre-formed, mature *P. aeruginosa* biofilms both qualitatively and quantitatively. Visual evidence of biofilm removal using CSLM confirmed significant reduction in biofilm thickness following exposure to PAAG (Figure 1) accompanied by increased bacterial killing within the deepest layers of the biofilm. Visualization also confirmed impaired biofilm growth at the level of the substratum, suggesting diffusion of PAAG through the biofilm layers. This observation was of particular significance as it suggested that PAAG has the potential to reduce biofilm integrity, potentially by dissolution and dispersal of the biofilm matrix (Figure 2).

The results of this study highlight the potential of PAAG to kill *P. aeruginosa* in biofilms, as well as disrupt its biofilm structure. Of significance, bactericidal effects were demonstrated using the laboratory isolate *P. aeruginosa* PAO1 as well as a mucoid CF clinical isolate, *P. aeruginosa* PA529. Visual evidence of biofilm removal by PAAG using CSLM demonstrated that biofilm thickness was significantly reduced following exposure to PAAG (Figure 1) and was accompanied by increased bacterial killing, including in the deepest layers of the biofilm. These studies also demonstrated bacterial killing and impaired biofilm growth at the level of the substratum, suggesting diffusion of the PAAG through the layer. This observation was of particular significance as it suggested that PAAG has the potential to reduce biofilm integrity, potentially by dissolution and dispersal of the biofilm matrix (Figure 2).

In the airways of patients with CF and other forms of bronchiectasis, biofilms are composed of a gel-like matrix of anionic extracellular polymeric substances including mucin, extracellular DNA and polysaccharides (12, 14). PAAG, a novel polycationic glycopolymer, has been demonstrated to normalize the structure of the airway mucin MUC5B in models of the CF airway. MUC5B linearization, a process that occurs normally in the setting of intact anion transport but is dysregulated in CF, contributes to abnormalities in mucus rheology and delayed mucociliary clearance (17). Given our previously published description on the effect of PAAG on the displacement of Ca^2+^ from MUC5B (17) and the known role of calcium in biofilm formation by *P. aeruginosa* we suspected and demonstrated that PAAG exhibited Ca^2+^ dependent anti-biofilm activity. To that end, we compared biofilm dissolution as a result of PAAG in varying levels of calcium. The presence of calcium was found to negate the effect of PAAG on biofilm dissolution, suggesting a similar mechanism of calcium displacement is necessary for PAAGs effects *on P. aeruginosa* biofilms.

In addition, the outer cell membranes of *P. aeruginosa* (and other Gram-negative bacteria) are dependent on Ca^2+^ for maintaining structural integrity interaction with negatively charged LPS on the outer membrane (30). Because of the impact of PAAG to displace Ca^2+^ in polymeric structures of mucins, we hypothesized that PAAG could also disrupt the cell membranes of *P. aeruginosa* isolates tested. Experiments exploring cell membrane disruption using a propidium iodide probe showed that PAAG rapidly permeabilized the *P. aeruginosa* cell membrane in a dose dependent manner. Furthermore, membrane depolarization studies demonstrated that PAAG caused rapid depolarization of the membrane as well. The effect of PAAG on the morphology of bacteria was also visualized using SEM imaging. SEM imaging provides visual evidence that exposure of planktonic *P. aeruginosa* cells to PAAG results in the deformation and disruption of the bacterial membrane.

Preventing and impairing bacterial adhesion to the epithelium is critical in preventing biofilm formation. Given the effects of PAAG on substratum viability and bacterial adhesion to A549 epithelial cells, in addition to anti-bacterial properties, we sought *in vivo* confirmation of reduced bacterial viability and colonization. Rats that received nebulized PAAG prior to and immediately following *P. aeruginosa* inoculation demonstrated decreased rates of *P. aeruginosa* colonization and reduced mean colony counts. Further, lung histology demonstrated that PAAG pretreatment resulted in decreased lung injury and neutrophilic inflammation. Given the prior findings that PAAG normalizes CF mucus rheology, this finding may be of additional clinical importance as neutrophilic inflammation is a key component in the vicious cycle of the development of bronchiectasis and chronic infection state (33).

*P. aeruginosa* is associated with the development of biofilm-associated infections in a variety of clinical scenarios including medical device infections and chronic infections including chronic airway colonization in CF, bronchiectasis, and other airway diseases. Novel mechanisms that result in prevention or dissolution of biofilms may alter the landscape of medical management of these infections. PAAG is a potential therapeutic which has previously been shown to normalize CF mucin structure and enhance mucociliary clearance. This study builds upon that finding by demonstrating potent antimicrobial and anti-biofilm effects of PAAG. Future studies evaluating the effect of PAAG on airway inflammation, in an environment in which *P. aeruginosa* infection is chronic, such as the CF lung, are warranted.

## Acknowledgments

We gratefully acknowledge Harvey-Mudd College and Dr. Tanenbaum for access and help with SEM imaging, and assistance from the UAB High Resolution Imaging and Microscopy Core Facility.

## Funding information

SMR: NIH R35 HL135816, NIH P30DK072482, CFF RDP R15RO.

WES: CFF SWORDS1810, CFF SWORDS 20G0

